# Identification of ASPDH as a novel NAADP-binding protein

**DOI:** 10.1101/2022.06.06.494947

**Authors:** Xiao He, Yunlu Kang, Lei Chen

**Author notes:** These authors contributed equally to this work (Xiao He, Yunlu Kang). Correspondence: Lei Chen.

## Abstract

NAADP is a signaling molecule that can induce calcium release from intracellular acidic stores. However, proteins that bind to NAADP are understudied. Here, we identify ASPDH as an NAADP-binding protein through biochemical purification from pig livers. ITC experiment using the recombinantly expressed protein shows a 1:1 binding stoichiometry and a Kd of 455 nM between NAADP and mouse ASPDH. In contrast, recombinantly expressed JPT2 and LSM12, two proteins previously identified as NAADP-receptors, show no binding in ITC experiments.

## Introduction

Calcium is an important second messenger that can initiate downstream signaling and has diverse physiological functions. Calcium signals arise from changes in local or global calcium ion concentration, which are mainly caused by calcium influx from plasma membrane or calcium release from intracellular calcium stores (1). Nicotinic acid adenine dinucleotide phosphate (NAADP) is a molecule that closely resembles nicotinamide adenine dinucleotide phosphate (NADP). NAADP could effectively immobilize calcium from acidic intracellular calcium stores to induce calcium signals (2–4). NAADP was commonly considered to be produced from the chemical reactions catalyzed by CD38 (5), while recent studies found that DUOX enzyme (6) and SARM1 (7) might also participate in NAADP synthesis in certain conditions. NAADP then binds to its receptor which further activates the TPC channel located on lysosome or endosome membrane to release calcium (8,9). Calcium signals evoked by NAADP have been found to mediate multiple physiological processes in mammals. For example, NAADP was shown to trigger calcium signals in pancreatic acinar cells (10), to promote pancreatic β cells to secret insulin under the stimulation of glucose (11), to induce the autophagy of hepatocytes when stimulated by LPS (12), and to depolarize the medullary neurons of rat (13). Moreover, NAADP signal is also implicated in several diseases, including T cell-mediated autoimmune diseases (14–17), cancers (18), and ventricular cardiomyocyte arrhythmias(19). Furthermore, NAADP signal plays roles in the infection process of certain viruses, such as ebolavirus (20) and MERS-CoV (21).

It is reported that the mammalian NAADP receptor binds NAADP with high affinity (Kd ∼5nM in mouse liver (9)). Radioactive ligand binding experiments have shown that NAADP-binding proteins are present in mouse livers and TPC2-overexpressed HEK293 cell line (9). In addition, photo-affinity labeling (PAL) experiments with radioactive compounds have identified multiple NAADP-binding proteins in mouse livers, pancreatic tissue, and cell lines including HEK 293, SKBR3, and Jurkat T-lymphocytes. These studies brought a 22-23 kDa protein into attention because its PAL signal could not be competed off by NADP (22–24). Recently, JPT2 (25,26) and LSM12 (27) were identified as the NAADP receptors through chemical biology approaches. Here, we described the identification of ASPDH from pig liver as a novel NAADP-binding protein through biochemical fractionation.

## Results

### Establishment of a non-radioactive NAADP-binding assay

To purify the NAADP-binding proteins, we sought to establish the NAADP-binding assay. Radioactive reagent ^32^P-NAADP was traditionally used for this purpose (9,28) but it is not commercially available, limiting its routine use in protein purification. Here, we develop a non-radioactive assay to detect the NAADP-binding ability of protein samples. It is reported that PEG-8000 could precipitate NAADP-binding protein from the sea urchin egg homogenate together with NAADP molecules bound on it (29). Moreover, NAADP could be detected by an established cycling assay with high sensitivity(30,31). We used a previously identified E98G mutant of *Aplysia* ADP-ribosyl cyclase to convert NAADP into NADP by base-exchange reaction at neutral pH (31,32). In the cycling assay, fluorescent resorufin is produced continuously along with the NADP cyclic reaction, and the rate of fluorescent increase is linear to the NADP concentration. Therefore, the amount of NAADP initially in the system was measured as the rate of fluorescence increased (Fig. 1A). Inspired by these advances, we first precipitate proteins with PEG-8000 to isolate the bound NAADP from the unbound NAADP (Fig. 1B) and then extract the bound NAADP from PEG-precipitated protein to determine its amount using the cycling assay.

**Figure 1.**
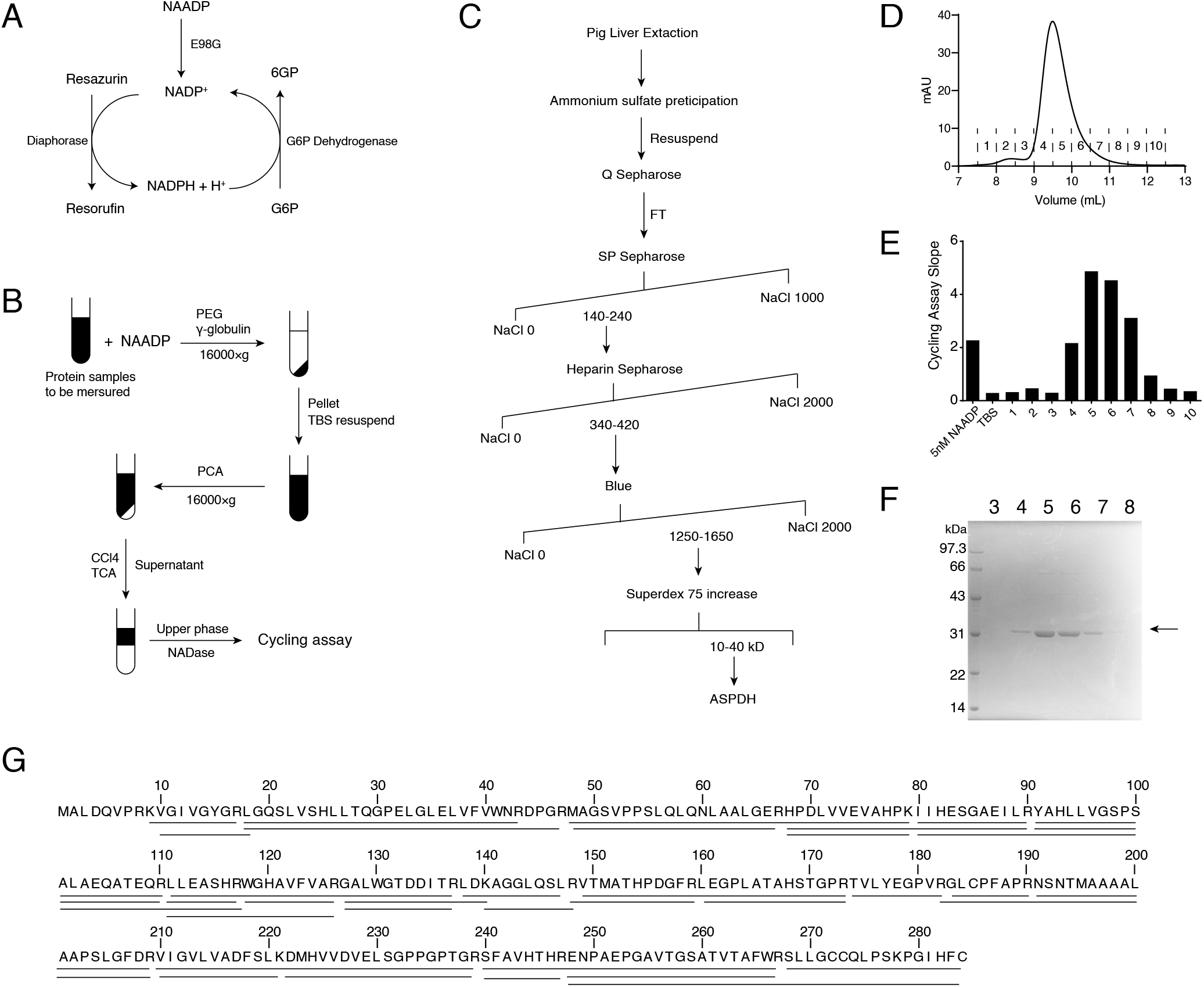
Biochemical purification of NAADP-binding protein from pig livers. (A) The reaction scheme of the cycling assay for NAADP concentration determination. NAADP was converted into NADP with the recombinant ADPRC E98G mutant. During the cycling assay, resazurin is converted into fluorescence resorufin with a reaction rate that is linear to the NADP concentration. (B) Isolation of protein-bound NAADP from free NAADP using PEG precipitation. Fractions containing NAADP are shown in black. (C) Schematic representation of the purification process of NAADP-binding proteins. Numbers represent the salt concentrations (mM) at which the active fractions were eluted. (D) The elusion profile of final size-exclusion chromatography. Fraction numbers were indicated. (E) NAADP-binding activity of fractions of the final size exclusion chromatography in (D). Bars represent the slopes of cycling assay and are proportional to the amount of NAADP that is bound to protein. (F) Coomassie blue-stained SDS-PAGE of fractions from the final size-exclusion chromatography. (G) The amino acid sequence of pig ASPDH. Peptides detected by LC-MS/MS were indicated by horizontal bars below.

### Purification of NAADP-binding proteins from pig liver

It was found that the liver tissue contains NAADP-binding protein and shows NAADP-induced calcium signals (12,22,24). We therefore used pig livers as the source for protein purification. The NAADP-binding protein was purified from the supernatant of pig liver homogenates after ultracentrifugation. Fractions with NAADP-binding activity were purified sequentially with ammonium sulfate precipitation and through several chromatographic columns, including anion-exchange chromatography (Q-sepharose), cation-exchange chromatography (SP-sepharose), affinity column (Heparin-sepharose and Blue-sepharose) (Fig 1C). After the final polishing step by the size-exclusion chromatography (Superdex 75 increase), the fractions with NAADP-binding activity contain a single protein band on SDS-PAGE with a molecular weight of around 30 kDa. (Fig 1 D-F). LC-MS/MS analysis identified the protein as aspartate dehydrogenase (ASPDH) with high coverage (Fig. 1G).

ASPDH is a member of amino acid dehydrogenases which catalyzes the dehydrogenation of L-aspartate to iminoaspartate (33), and uses <u>nicotinamide adenine dinucleotide</u> (NAD) or NADP as the electron acceptor. It is reported that ASPDH is involved in NAD biosynthesis in some anaerobic hyperthermophiles. In addition, ASPDH may catabolize aspartate to provide energy and nitrogen in some mesophilic bacteria strains (34). However, the function of eukaryotic ASPDH is unknown. The estimated molecular weight of pig ASPDH is 31 kDa, which agrees well with the migration pattern of the active fractions on SDS-PAGE.

### Recombinant ASPDH can bind NAADP

To further confirm that ASPDH can bind NAADP, we expressed His-tagged mouse ASPDH (His-mASPDH) in *E*.*Coli* and purified the protein to apparent homogeneity through Ni-NTA, ion-exchange chromatography and size-exclusion chromatography. The migration of recombinant His-mASPDH on SDS-PAGE (Fig. 2A) and its peak position on size-exclusion chromatography (Fig. S2A) match the active factions purified from pig liver (Fig. 1D-F). To quantify the thermodynamics of NAADP-binding to ASPDH, we performed ITC measurements. We found that ASPDH binds NAADP with a 0.78±0.049 stoichiometry and with a Kd of around 455.4±230.6 nM. We also found that ASPDH binds NADP with a stoichiometry of 0.77±0.096, while with a Kd of 958.2±202.9 nM (Fig. 2 B-D). The binding stoichiometry of NAADP or NADP over ASPDH is lower than 1:1, possibly due to the over-estimation of protein concentration by the Bradford assay.

**Figure 2.**
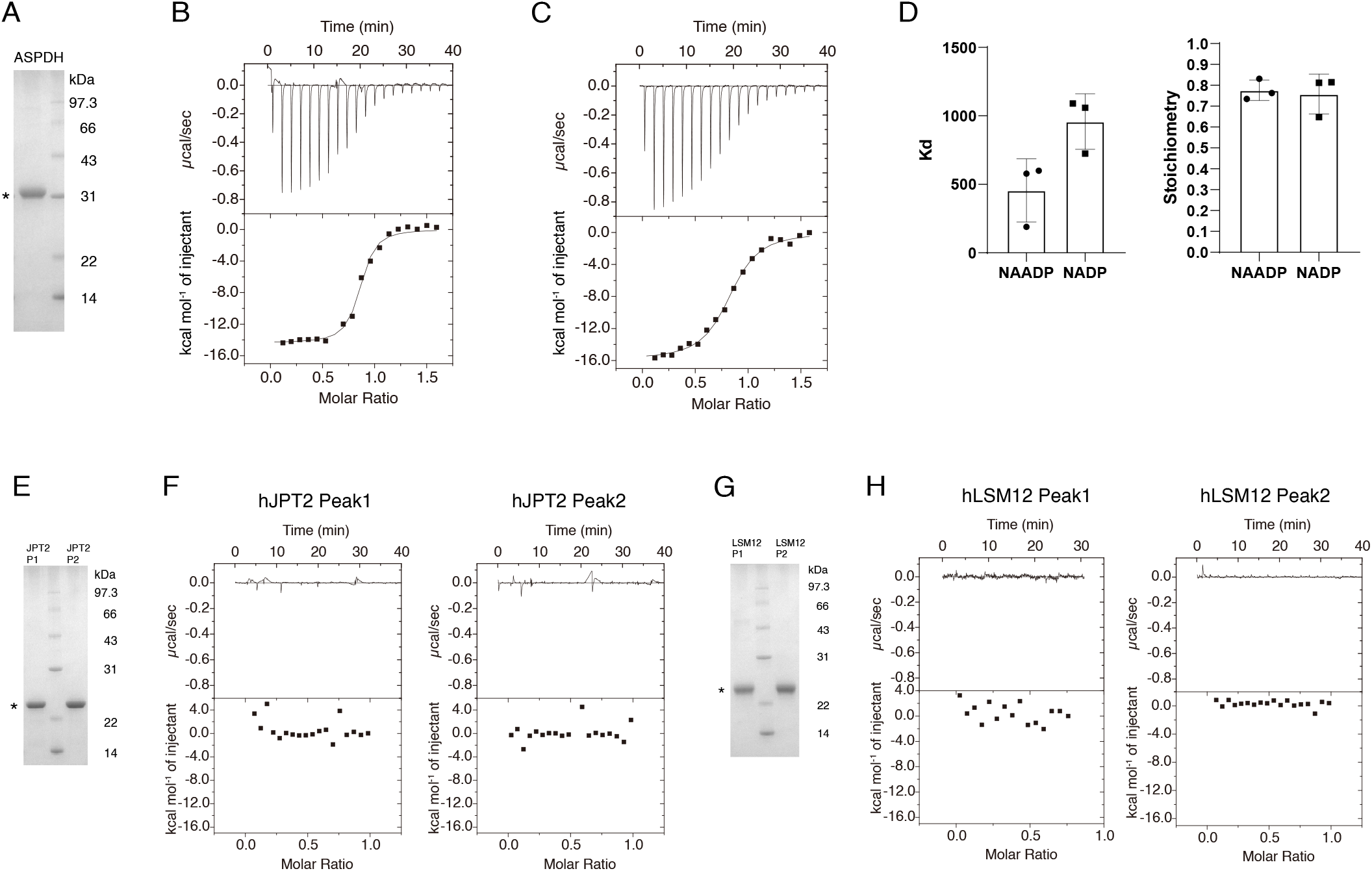
Thermodynamics of NAADP-binding process measured with ITC. (A) Coomassie blue-stained SDS-PAGE of His-mASPDH used in ITC. (B) Representative ITC measurement shows that His-mASPDH was able to bind NAADP. Integrated data (lower panel) were obtained by fitting raw data (upper panel) with the one-site model. (C) Representative ITC measurement shows that His-mASPDH was able to bind NADP. Integrated data (lower panel) were obtained by fitting raw data (upper panel) with the one-site model. (D) Statistic results of Kd and stoichiometry of NAADP and NADP binding to ASPDH. The Kd and stoichiometry for NAADP are 455.4±230.6 nM and 0.78±0.049. The Kd and stoichiometry for NADP are 958.2±202.9 nM and 0.77±0.096. Results were obtained from three independent ITC experiments. Data were presented as mean ± standard deviation. (E) Coomassie blue-stained SDS-PAGE of hJPT2 used in ITC. (F) ITC measurements show that hJPT2 proteins from both peaks of size-exclusion chromatography do not bind NAADP. The experiment has been performed for twice with similar result. (G) Coomassie blue-stained SDS-PAGE of hLSM12-His used in ITC. (H) ITC measurements show that hLSM12-His proteins from both peaks of size-exclusion chromatography do not bind NAADP. The experiment has been performed for twice with similar result.

### Recombinant LSM12 and JPT2 do not bind NAADP

LSM12 and JPT2 were two proteins that were recently identified to be the NAADP receptors in several cell lines (25–27). To quantify the thermodynamics of the binding between NAADP and LSM12 or JPT2 and to compare them with ASPDH, we recombinantly expressed these two proteins in *E*.*Coli* (Fig. S1) and purified them to apparent homogeneity for ITC experiments (Fig. 2E-H). Recombinant human LSM12 was expressed with a C-terminal His-tag (hLSM12-His). Human JPT2 was expressed with an N-terminal GST-tag (GST-hJPT2), and the GST tag was cleaved and removed before ITC experiments. Both proteins were eluted as two peaks on size-exclusion chromatography (Fig. S2B, C). We collected these two peaks separately for the ITC measurement. To our surprise, we could not detect any changes when NAADP was titrated into hLSM12-His or hJPT2 protein using ITC, suggesting no binding between NAADP and LSM12 or JPT2 (Fig. 2E, G).

## Discussions

In this work, we have isolated ASPDH from pig liver tissue and demonstrated that it is a novel NAADP-binding protein. It is reported that ASPDH might use NAD or NADP as its substrate. Moreover, structures of ASPDH from *Thermotoga maritima* and *Archaeoglobus fulgidus* have revealed that ASPDH has a Rossmann fold domain that binds NAD (33,35) (Fig. S3A, B). The structure of eukaryotic ASPDH, such as mouse ASPDH, predicted using AlphaFold2 (36) also shows a conserved Rossmann fold domain (Fig. S3C, D). Given the high structural similarity between NADP and NAADP, we speculate the Rossmann fold domain of ASPDH could bind NAADP as well. Our ITC experiments show ASPDH binds NAADP with medium affinity (∼455.4 nM, Fig. 2B), which does not match the high affinity of NAADP receptor (Kd∼5nM (9)). Moreover, ASPDH can also bind NADP with similar affinity (∼958.2 nM, Fig. 2C), suggesting ASPDH has low selectivity between NAADP and NADP. Furthermore, the ASPDH gene is transcribed most abundantly in livers and kidneys in humans, while NAADP signal is more broadly distributed (37). The medium NAADP affinity, low selectivity between NAADP and NADP, and limited tissue distribution of ASPDH suggest it is not the bona fide NAADP receptor and its biological function in NAADP signaling remains elusive.

It is reported that recombinant LSM12 and JPT2 bind NAADP with high affinity (the Kd of both proteins were around 20 nM) and higher selectivity in preference to NADP using competitive radioligand binding assay (25–27). Moreover, LSM12 might have an additional low affinity site that binds NAADP with a Kd of about 5 μM (27). However, the predicted structures of hLSM12 and hJPT2 (36) do not have any recognizable domain that can bind NAADP or NADP (Fig. S3E, F). Together with our ITC experiments, these data question the NAADP-binding capability of hLSM12 and hJPT2, the only two proposed NAADP-receptors.

## Figure Legends

**Figure S1.**
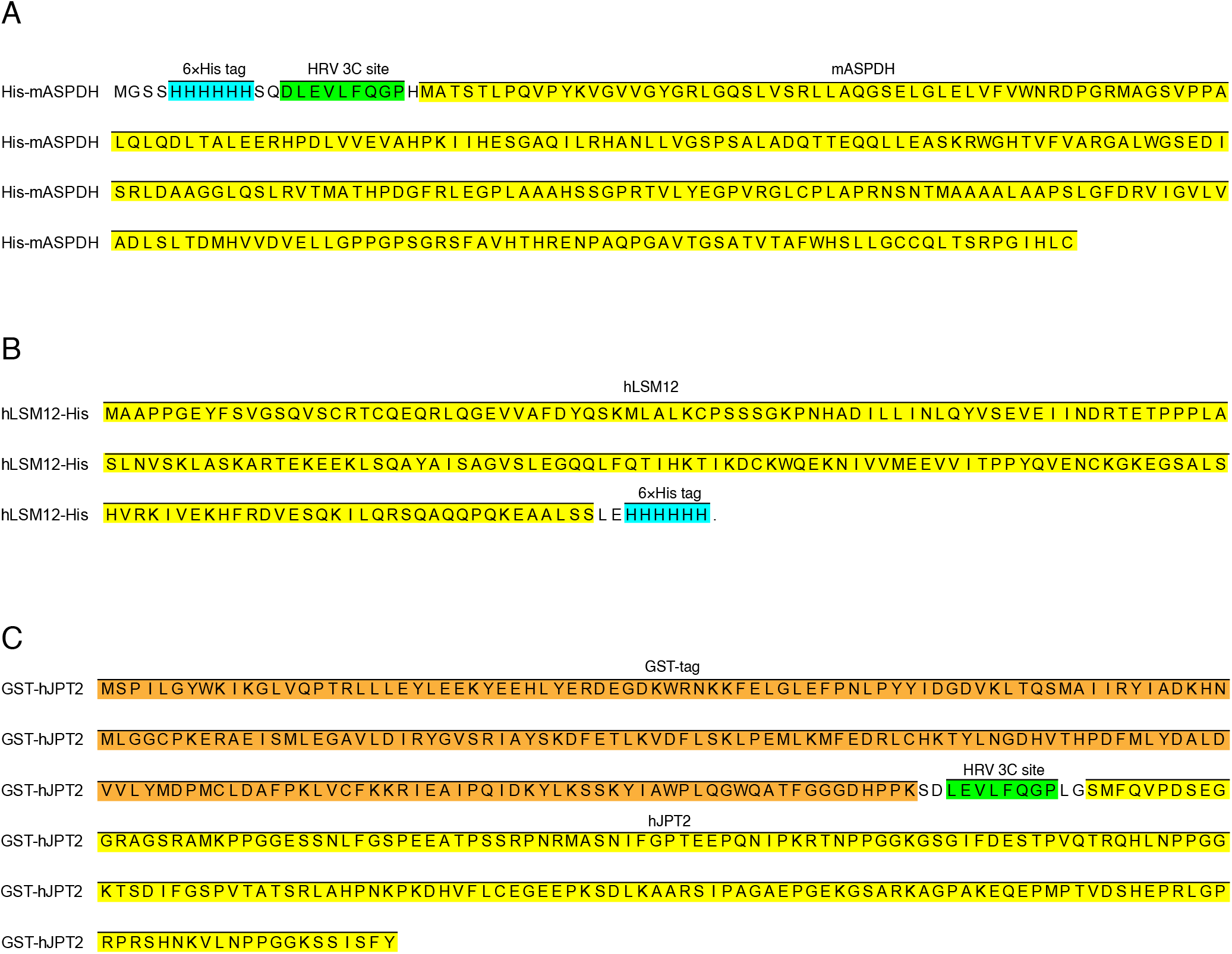
Sequences of recombinant proteins were used in this study. His tags were labeled with blue, HRV 3C sites were labeled with green, GST tag was indicated with orange and the sequences of the target proteins were indicated with yellow. (A) The sequence of recombinantly expressed His-mASPDH. (B) The sequence of recombinantly expressed hLSM12-His. (C) The sequence of recombinantly expressed GST-hJPT2

**Figure S2.**
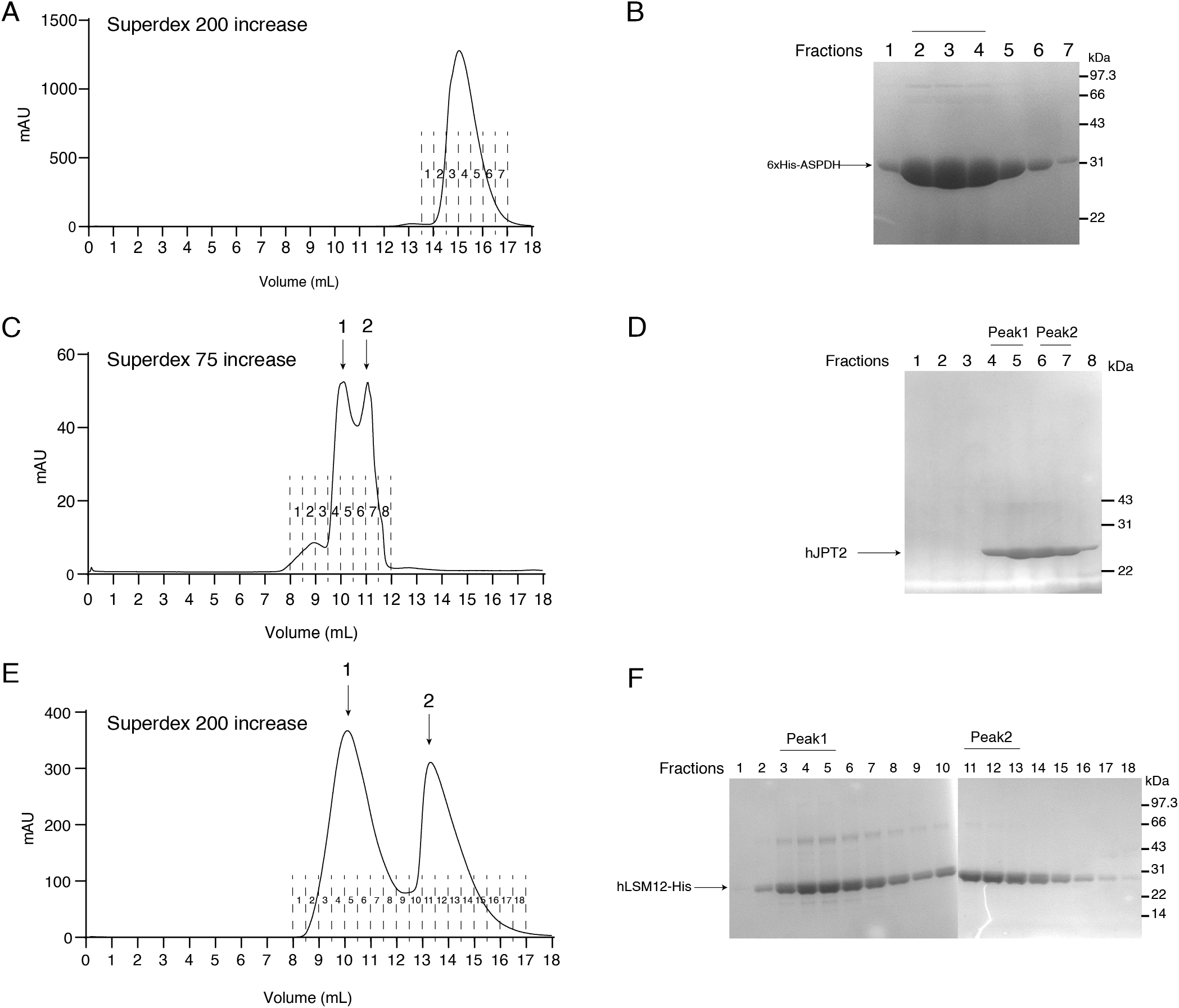
The elution profiles of target protein on Size-exclusion chromatography (SEC). (A) SEC profile of His-mASPDH. The column and fraction numbers were indicated. (B) Coomassie blue-stained SDS-PAGE of SEC fractions of His-mASPDH. The His-mASPDH was indicated with an arrow. Fractions 2-4 were collected for ITC experiments. (C) SEC profile of hJPT2 whose GST tag had been removed. The column and fraction numbers were indicated. Individual peaks were marked with arrows. (D) Coomassie blue-stained SDS-PAGE of SEC fractions of hJPT2. The hJPT2 was indicated with an arrow. Fractions 4-5 and 6-7 were collected respectively as Peak1 and Peak2 for ITC experiments. (E) SEC profile of hLSM12-His. The column and fraction numbers were indicated. Individual peaks were marked with arrows. (F) Coomassie blue-stained SDS-PAGE of SEC fractions of hLSM12-His. The hLSM12-His was indicated with an arrow. Fractions 4-6 and 11-13 were collected respectively as Peak1 and Peak2 for ITC experiments.

**Figure S3.**
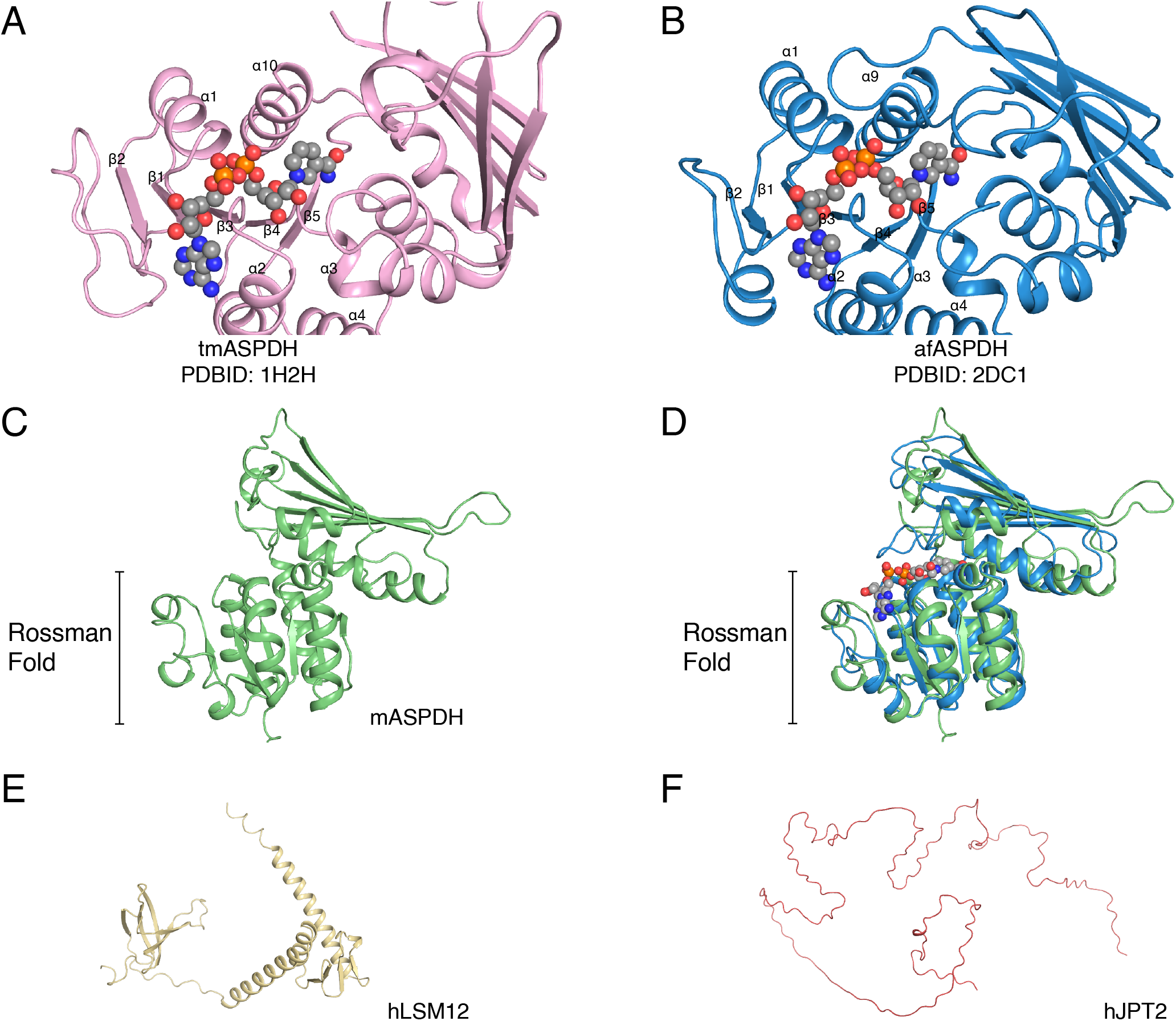
Rossman fold domain of ASPDH might bind NAADP. (A) Structure of *Thermotoga maritima* ASPDH (tmASPDH, PDB ID: 1H2H), showing the Rossman Fold domain with NAD molecule bound on it. (B) Structure of *Archaeoglobus fulgidu* ASPDH (afASPDH, PDB ID: 2DC1), showing the Rossman Fold domain with NAD molecule bound on it. (C) Structure of mASPDH predicted with AlphaFold2. (D) Structural alignment of afASPDH (violet) and predicted mASPDH (green). The structures of Rossman Fold domains between the two proteins were highly conserved. The NAD molecule bound in afASPDH was shown to indicate the position of the dinucleotide in the whole structure. (E) Structure of hLSM12 predicted with AlphaFold2. (F) Structure of hJPT2 predicted with AlphaFold2.

## Experimental procedures

### Materials

NAADP was purchased from Tocris Bioscience. NADP was purchased from Abcam. γ-globulin, PEG-8000, Flavin mononucleotide (FMN), resazurin, Glucose-6-phosphate (G6P), G6P dehydrogenase (G6PDH) and Diaphorase were purchased from

Sigma. Phosphatase inhibitor cocktail A was purchased from Beyotime.

### Purification of enzymes

*Aplysia* ADP-ribosyl cyclase (ADPRC) and NADase were cloned into pPICZα and the E98G mutant on ADPRC was generated by Quickchange PCR. ADPRC with E98G mutant (E98G) and NADase were expressed in X-33 *Pichia pastoris* and purified with affinity chromatography and ion-exchange chromatography. E98G, NADase and all commercially available enzymes used in NAADP cycling assay were further treated with activated charcoal to remove the contamination of NADP and its derivates.

### Isolation of bound NAADP from free NAADP

100 μL protein samples were mixed with 100 μL 2× Glu-IM Buffer (500 mM potassium gluconate, 500 mM N-methylglucamine, 40 mM HEPES, 2 mM MgCl_2_, pH 7.2, treated with activate charcoal) and added with protease inhibitor (5 mM ethylenediaminetetraacetic acid (EDTA), 2 mM ethylene glycol ltetraacetic acid (EGTA), 1 μg/ml aprotinin, 1 μg/ml pepstatin, 1 μg/ml leupeptin, 1 mM phenylmethanesulfonyl fluoride (PMSF)) and phosphatase inhibitor (5 mM sodium fluoride, 1 mM sodium pyrophosphate, 1 mM β-glycerophosphate and 1 mM sodium orthovanadate). The mixture was incubated with 100 nM NAADP on ice for 40 min. 4 mg/mL γ-globulin and 15% PEG 8000 in final concentration was added to protein samples. The mixture was incubated on ice for 30 min and centrifuged at 16, 000 ×g. After being washed once with 15% PEG 8000, precipitates were resuspended with TBS (20 mM Tris-HCl, 150 mM NaCl) and NAADP was released by adding <u>perchloric acid</u> (PCA) to 0.6 M in final concentration. Denatured proteins were removed by centrifugation and supernatant containing released NAADP was extracted with 25% Chloroform: Thionylamine (3:1 v/v) solution to remove the excessed PCA. The aqueous phase was isolated by centrifugation at 1, 000 ×g for 5 min, and the upper layer was collected, added with 4 mM MgCl_2_, 10 mM Tris-HCl (pH = 7.5) and NADase to eliminate NADP by digestion under 37 °C overnight.

### NAADP cycling assay

The 2× reaction buffer contained 20 μM FMN, 20 μM resazurin, 2 mM G6P, 100 mM nicotinamide, 100 mM sodium phosphate (pH 8), 0.2 U/ml G6PD, 20 μg/mL diaphorase with or without 93 μg/ml E98G protein. Samples treated with NADase overnight were boiled at 95 °C for 20 min to inactivate NADase, and then the denatured NADase was removed by centrifugation (16, 000 ×g, 10 min). The supernatant was mixed with equal volume of 2× reaction buffer and incubated at room temperature for 1 h. The velocity of fluorescence increase was performed by measuring the fluorescence every 1 minute. A reaction system without E98G protein was used as the negative control.

### Preparation of pig liver extraction

Pig livers purchased at the local butcher’s shop were cut into 1 cm^3^ cubes and homogenized in Buffer A (50 mM Tris (pH 7.5), 150 mM NaCl, 5 mM EDTA, 2 mM EGTA and 1 mM PMSF) with a disperser (IKA T18 digital ULTRA-TURRAX). The crude lysis was ultracentrifugated at 35, 000 rpm for 40 min at 4°C (Type 45Ti). The supernatant was carefully transferred with pipette for further purification.

### NAADP binding proteins purification from pig liver extraction

NAADP binding proteins in pig liver extraction was salted out with 3.5 M amino sulfate (pH 8.0) on ice for 30 min and the final proportion for amino sulfate was 35% (v/v). Precipitated proteins were isolated by centrifugation, dissolved in Buffer B (50 mM Tris (pH 7.5), 50 mM NaCl, 5 mM EDTA, 2 mM EGTA and 1 mM PMSF), and centrifuged at 16, 000 ×g (JA14.4) for 10 min. The supernatant was added with 15% (v/v) ethanol on ice for 30min, and centrifuged at 16, 000 ×g for 20 min. The supernatant was diluted with H_2_O until the conductance was lower than 5000 μS, then subjected to anion-exchange chromatography (Q FF, CYTIVA). The flow-through of Q column was adjusted to pH 5.91 using 3 M sodium acetate (pH=5), and diluted with H_2_O until the conduction was lower than 5000 μS. Diluted flow-through was loaded onto cation-exchange chromatography (SP 5 mL HP, CYTIVA), and eluted with a linear concentration gradient of NaCl from 0-1 M for 25 mL and 1 M for another 10 mL in 20 mM MES (pH = 6). The binding active fractions were pooled and adjected to pH=8 with 2 M Tris-HCl (pH=8), then diluted with H_2_O until the conductance was lower than 4000 μS. Heparin Sepharose was then performed in series to further purify the NAADP-binding proteins, and elution was performed with 0-1 M NaCl for 25 mL and 2 M NaCl for another 10 mL. Notably, the NAADP-binding activity was separated into two peaks on Heparin Sepharose and only proteins in the second peak could be purified further and identified by LC-MS/MS. The fractions in the second activity peak were combined, diluted and loaded onto Blue Sepharose column. Proteins with activity were eluted using 0-2 M NaCl for 40 mL and 2 M NaCl for another 10 mL. For final polishing, fractions showing NAADP-binding activity were pooled and loaded onto size-exclusion chromatography (Superdex 75 increase, CYTIVA) in TBS buffer.

### SDS-PAGE of protein samples

The protein samples were mixed with 6× SDS Loading Buffer containing 60% glycerol (v/v), 300 mM Tris-HCl (pH 6.8), 12% SDS (w/v), 0.6% bromophenol blue (w/v), and separated on 12% polyacrylamide gels by performing SDS-PAGE. Gels were stained with 0.25% Coomassie blue (R-250) in 50% ethanol and 10% acetic acid. Bands in the stained gel were carefully cut off with a scalpel for MS analysis.

### Expression and purification of recombinant proteins

We cloned mouse ASPDH (mASPDH) into a modified pET expression vector with N-terminal His-tag (pPH) and human LSM12 (hLSM12) into a pET expression vector with C-terminal His-tag. We cloned human JPT2 (hJPT2) into a pGEX expression vector with N-terminal GST-tag. Recombinant proteins were all expressed in NiCo21 *E*.*Coli* strain, and cells were harvested after being induced with IPTG at 16°C for 20 h. After lysed with sonication and centrifugation, the supernatant was subjected to affinity chromatography (for His-mASPDH, Talon; for hLSM12-His, Ni-NTA; for GST-hJPT2, GST). His-tagged ASPDH and LSM12 were subjected to ion-exchange chromatography (Q HP, CYTIVA) and eluted with an increasing concentration of NaCl (0-1 M) in 20 mM Tris-HCl (pH = 7.5). For GST-hJPT2, the GST tag was removed by ppase digestion, after which the pH was adjusted to 6.0 with 1 M MES (pH = 5.9) and the free GST tag and ppase were removed with cation-exchange chromatography (SP HP). hJPT2 was eluted with an increasing concentration of NaCl (0-1 M) in 20 mM MES (pH = 5.9). Size-exclusion chromatography (for His-mASPDH and hLSM12-His, Superdex 200 increase, and for hJPT2, Superdex 75 increase) were performed to polish the purification of recombinant proteins and to substitute their buffers with ITC buffer, which contained 20 mM Tris-HCl, 150 mM NaCl and 2 mM tris (2-carboxyethyl) phosphine (TCEP).

### ITC measurement

The protein concentration of His-mASPDH was measured with Bradford Assay Kit (Beyotime). The protein concentrations of hLSM12-His and hJPT2 were estimated by measuring their UV absorbance at 280 nm and calculated with their estimated molar absorption coefficient (13,575 for hLSM12-His, and 1,615 for hJPT2). Concentrations of NAADP and NADP were determined by measuring UV absorbance at 254 nm, with a molar absorption coefficient of 18300. 3.4 mM NAADP and 24.5 mM NAADP in water, which were adjusted to pH=7 with NaOH and HCl, were diluted to proper concentrations with ITC buffer, and so were the proteins. During ITC experiment, ligands were loaded stepwise into 300 μL proteins on MicroCal ITC200 instrument. The first drop contained 1 μL while the following drops contained 2 μL each, and 19 drops were loaded in total. Data were processed with Origin 7.0. For His-mASPDH, integrated data were obtained by fitting raw data with the one-site model. After titration, the proteins were collected and subjected to SDS-PAGE.

## Data availability

All data are contained within this manuscript.

## Acknowledgement

We thank Prof. Yongjuan Zhao for providing the cDNA of *Aplysia* ADP-ribosyl cyclase. We thank the Protein Chemistry Facility Center, School of Life Sciences, Tsinghua University and National Centre of Protein Sciences at Peking University for LC-MS/MS assay. We thank National Centre of Protein Sciences at Peking University for ITC experiments.

## Author contributions

L.C. conceived the project. Y.K. established the NAADP-binding assay. X.H. and Y.K. purified the ASPDH from pig livers. X.H. performed the ITC experiments. All authors contribute to the manuscript preparation.

## Funding and additional information

The work is supported by grants from the National Natural Science Foundation of China (91957201, 31870833 and 31821091 to L.C.) and Center For Life Sciences (CLS). Y.K. is supported by the Boya Postdoctoral Fellowship.

